# Advancing species identification: A non-invasive molecular approach through spider silk proteome analysis

**DOI:** 10.1101/2024.05.09.593458

**Authors:** Phillip K Yamamoto, Keizo Takasuka, Masaru Mori, Takeshi Masuda, Nobuaki Kono

**Affiliations:** Systems Biology Program, Graduate School of Media and Governance, Keio University, Fujisawa, Kanagawa, 252-0882, Japan; Institute for Advanced Biosciences, Keio University, Tsuruoka, Yamagata, 997-0017, Japan; Faculty of Environmental and Information Studies, Keio University, Fujisawa, Kanagawa, 252-0882, Japan

**Keywords:** Spider silk, Species identification, Proteome analysis, Protein

## Abstract

Species identification is crucial in various scientific disciplines such as biology, ecology, medicine, and agriculture. While traditional methods rely on morphological characteristics, DNA barcoding has gained popularity due to its molecular biology approach. Nonetheless, DNA barcoding can be problematic for small animals such as insects, as it requires damaging their bodies for DNA extraction, impacting subsequent breeding and experiments. In this paper, we propose a non-invasive molecular method for species identification that examines the protein composition of animal produced biomaterials. We chose spider silk, with species-specific protein sequences, as our subject of analysis. First, we established a universal silk-dissolving method that applies to silks from various species. We constructed a bioinformatics pipeline employing metrics of significant difference through proteomic analysis to identify spider species by analyzing peptide sequences present in silk proteins. As a result, we achieved a species identification accuracy of 86% across15 species. An appropriate reference dataset was successfully created, in addition, we also discovered some species are difficult to distinguish due to sequence similarities. This technology has been confirmed to be applicable to spider webs taken from the field. This non-invasive approach can complement DNA barcoding, especially in situations where it is infeasible, such as in studies involving spider-parasitoid wasps that eat spiders. Furthermore, it can be applied to other organisms that release biological substances, such as silkworm pupae, termite digestive enzymes, and tick saliva, aiding in species identification and pest control efforts.

## INTRODUCTION

Species identification of organisms is an important basic technique not only in biological and ecological research (Elphick, 2008; Farnsworth et al., 2013; Kürzel et al., 2022) but also in pathogen identification in medicine (Becker et al., 2003; Lou et al., 2010) and pest identification in agriculture (Ashfaq & Hebert, 2016). Traditionally, species identification has been based on morphological characteristics which required special knowledge of the organism (Trail, 2021), and this approach was the predominant method until about 20 years ago. There are problems with misidentification due to phenotypic plasticity and lack of knowledge. Additionally, identification of many individuals poses a challenge as morphological characteristics are only evident during a particular stage of life or in a certain sex (Ajmal et al., 2014). As analyzing technologies has advanced, there has been a growing emphasis on molecular information as a more precise identifier than morphological characteristics. One key method they use is called DNA barcoding (Hebert, et al., 2003a; Hebert, et al., 2003b). This technique involves taking a body part of an organism, sequencing the DNA molecules contained in it through DNA amplification, and comparing it to databases with existing biological DNA sequences for identification. As of today, DNA barcoding is commonly used because it can be performed by anyone with the equipment and does not require any special knowledge (Ankola et al., 2021).

However, DNA barcoding is not suitable for studies necessitating continuous observation and experimentation, as it is invasive to living organisms. This is especially true for identifying smaller organisms such as insects and spiders, where any damage inflicted can adversely affect their growth. For instance, in long-term studies observing the growth of living organisms, DNA barcoding, can only be performed after all observations have been completed. Additionally, DNA barcoding is inapplicable in the absence of the target organism. For example, employing DNA barcoding is difficult in ecological studies of spider-parasitic wasps; in the initial discovery that *Eriostethus rufus* might be parasitic on *Trichonephila clavata* (Joro spider), the spider was already killed by the wasp with the host remnant gone away. Therefore, the host spider was assumed to be *T. clavata* based on the shape of the spider web and the species of spider that was located nearby (Takasuka, 2021). Without the main body of the organism, invasive DNA barcoding could not be employed, and important discoveries could be missed.

Thus, we propose a non-invasive molecular method for species identification that examines the protein composition of animal produced biomaterials. Here, we chose spider silk, with species-specific protein sequences, as our subject of analysis. Recently, the spider silk gene sequence data are increasing due to the extensive organization of genome information (K. Arakawa et al., 2022; Babb et al., 2017; Berger et al., 2021; Kono et al., 2019, 2020; Kono, Nakamura, & Arakawa, 2021; Kono, Nakamura, Mori, et al., 2021; Kono, Ohtoshi, et al., 2021) and transcriptome information (Arakawa et al., 2022). Currently, there are more than 1, 000 species of spider silk genes that have been identified and a database necessary for proteome analysis has recently emerged. Additionally, proteome databases have been progressively organized (Okuda et al., 2017) and there are more direct assessments of protein expression in biomaterials, beyond DNA and RNA level (Kazuharu Arakawa et al., 2021; Kono, Nakamura, Ohtoshi, et al., 2021). Hence, our research aims to develop a method for species identification from spider silk through proteome analysis.

## MATERIALS AND METHODS

### 2.1 Sampling and rearing

Adult females of *Dictyna felis* (family Dictynidae), *Parasteatoda tepidariorum* (family Theridiidae), *Cyclosa laticauda* (family Araneidae), *Cyclosa octotuberculata* (family Araneidae), *Yaginumia sia* (family Araneidae), *Lycosa suzukii* (family Lycosidae) were collected from Yamagata Prefecture, Japan (April 2023). Petri dishes with a diameter of 10 cm and a height of 3 cm, sterilized by UV light, were used for rearing. Spiders were individually reared at 20°C for two weeks while weaving. No feeding was performed. The identification of spider specimens was performed based on morphological characteristics and sequence identification of cytochrome *c* oxidase subunit 1 (*COX1*) in the Barcode of Life Data System (BOLD: http://barcodinglife.org).

### 2.2 Silk collection

Silks were collected with a needle manually from the spider web, and an electric drill was used directly for each spider. A minimum of 0.01 mg was sampled from each individual. The collected spider silks were stored at room temperature until used.

### 2.3 Protein extraction

Collected spider silk was immersed in the lysis buffer [9M LiBr or 8M Gdn-HCl] and boiled at 95°C for 20 min followed by cooling at room temperature for 10 min. After mixing, the sample was sonicated using a Biorupter Ⅱ (BM Equipment Co., Ltd.) with high energy for 20 min at intervals of 1 min. This process was repeated twice. After the first lysis, the boiling time at 95℃ was shortened to 10 min. The Pierce™ BCA Protein Assay Kit (Thermo Fisher Scientific) was used to quantify the amount of extracted protein.

### 2.4 LC-MS/MS for DDA using timsTOFPro

The lysate was treated with dithiothreitol at 37°C for 30 min followed by iodoacetamide at 37°C for 30 min in the dark. Following the five- or ten-fold dilution (8M Gdn-HCl or 9M LiBr) with 50 mM ammonium bicarbonate, the sample was incubated with sequence grade chymotrypsin (Promega) at 37°C for 18 hours. The digests were acidified with trifluoroacetic acid and desalted using a home-made C18-StageTip (a 200 µL micro pipet tip packed with a Empore SDB-XC disk [GL Science] and 10 mg of YMC GEL ODS-AQ particles [YMC CO., LTD]). The Stage Tip was briefly washed with 0.1% trifluoroacetic acid (TFA) 80% acetonitrile (ACN) and then equilibrated with 0.1% TFA 2% ACN. The sample solution was loaded followed by washing with 0.1% TFA 2% ACN. The retained peptides were eluted with 0.1% TFA 80% ACN. The eluate was dried under reduced pressure and dissolved in 0.1% formic acid (FA) 2% ACN for LC-MS/MS.

Nano liquid chromatography tandem mass spectrometry (nanoLC–MS/MS) was performed using a nanoElute ultrahigh-performance LC apparatus and a timsTOFPro mass spectrometer (Bruker Daltonics, Bremen). Each digested sample was injected into a self-packed needle column (ACQUITY UPLC BEH C18, 1.7 μm, Waters, Milford, MA, 75 μm i.d. × 250 mm) and separated by linear gradient elution with two mobile phases, A (0.1% formic acid in water) and B (0.1% formic acid in acetonitrile), at a flow rate of 280 μL/min. The composition of mobile phase B was increased from 2 to 35% in 100 min, changed from 35 to 80% in 10 min, and kept at 80% for 10 min. The separated peptides were ionized at 1600 V and analyzed by parallel accumulation serial fragmentation scan. Precursor ions were selected from the 12 most intense ions in a survey scan (precursor ion charge: 0–5, intensity threshold: 1,250, and target intensity: 10, 000) with ddaPASEF mode (Meier et al., 2018).

### 2.5 LC-MS/MS for DIA using Orbitrap Exploris 480

Sample preparation for a liquid chromatography tandem mass spectrometry (LC-MS/MS) was performed with label-free method. For desalting, a Stage Tip (Rappsilber et al., 2003) was prepared by filling two sheets of SDB-XC into the tip of a 200 μL tip. 200 μL of protein elution solution [0.1% TFA 80% ACN] was added and centrifuged at 1, 000 × g for 2 min. 200 μL of washing solution [0.1% TFA 5% ACN] was added and centrifuged at 1, 000 × g for 2 min. The protein concentration was adjusted with 9M LiBr to 5 μg/20 μL per sample. 2 μL of reducing solution [100 mM DTT 50 mM NH4HCO3] was added to the samples and incubated at 37°C for 30 mins, followed by the addition of 2 μL of alkylating solution [500 mM IAA 50 mM NH4HCO3] and incubation at 37°C for 30 min in the dark. After incubation, 180 μL of 50 mM NH4HCO3 and 1 μL of protein fragmentation solution [0.5 μg/μL Chymotrypsin 1 mM HCl] were added and incubated at 37°C for 18 hours. After adding 2 μL of TFA, it was poured into a Stage Tip for 200 μL and centrifuged at 1,000 × g for 3 min. Peptides were purified by adding 200 μL of washing solution and centrifuging at 1,000 × g for 3 min, followed by adding 50 μL of protein eluent and centrifuging at 1,000 × g for 3 min, repeated twice. The purified peptides were placed in a 96-well plate, dried in a vacuum concentrator for 2 hours, and 7.5 μL of eluent [0.1% TFA] was added.

Vanquish Neo UHPLC system (Thermo Fisher Scientific) and Orbitrap Exploris 480 (Thermo Fisher Scientific) were used. The LC was performed with a C18 packed analytical column (75 μm ID, 12.5 cm length, Nikkyo Technos). The mobile phase consisted of (A) 0.1% formic acid and (B) 0.1% formic acid in 80% acetonitrile. A linear gradient was performed as follows: 0 min %B: 5%, 50 min %B: 15%, 120 %B: 40%, 121 min %B: 100%, 130 min %B: 100%. The parameters were follows: MS data acquisition method: DIA, Method Duration = 125 min, Ion Source Type = NSI, Spray Voltage: Positive Ion = 2,300 V, Ion Transfer Tube Temp = 275 °C, Internal Mass Calibration = EASY-ICTM, Application Mode = Peptide, Expected Peak Width = 30 s, Default Charge State = 3, Advanced Peak Determination = True, MS1 Orbitrap Resolution 60000, MS1 Scan Range = 500-740 m/z, Polarity = Positive, DIA Window Mode = Center Mass, HCD Collision Energies 22, 26, 30 %, MS2 scan range 200-1800 m/z, MS2 resolution: 30,000.

### 2.6 Proteome data analysis

PEAKS Studio Xpro ver20200219 (Zhang et al., 2012) was used to analyze the mass spectra obtained from the proteome analysis. The following options were used for the analysis. Parent Mass Error Tolerance: 20.0 ppm, Fragment Mass Error Tolerance: 0.05 Da, Precursor Mass Search Type: monoisotopic, Enzyme: Chymotrypsin, Max Missed Cleavages: 2, Digest Mode: Specific, Fixed Modifications: Carbamidomethylation: 57.02, Variable Modifications: Acetylation (Protein N-term): 42.01, Oxidation (M): 15.99, Max Variable PTM Per Peptide: 3, Taxon: All, Contaminant Database: contami-nants_MaxQuant1.6.10.43 (no perfect match), Searched Entry: 2,069,477, FDR Estimation: Enabled, Merge Options: no merge, Precursor Options: corrected, Charge Options: no correction, Filter Charge: 1 – 5, Process: true, Associate chimera: yes. The analysis was also performed using DIA-NN (1.8.1) (Demichev et al., 2020). Default settings have been used without match between runs feature, allowing two chymotryptic missed cleavages.

The reference sequence list used 11,155 spider silk gene sequences of 1,098 spider species from Spider Silkome Database (https://spider-silkome.org/) (Arakawa et al., 2022), excluding duplicated sequences and sequences annotated as “Spidroin” and “Ampullate spidroin”.

### 2.7 Datasets

Silk proteome analysis data, including JPST001986, JPST001179, JPST001094, and JPST001096, were sourced from jPOSTrepo (Okuda et al., 2017).

### 2.8 Bioinformatics analysis

All bioinformatics analyses were performed using Python custom scripts. The statistical analyses and visualizations were implemented using the R package (v 4.0.5). The sequence alignment was produced by MAFFT (v7.505) (Katoh & Standley, 2013) using the default parameters and the alignment was visualized with Jalview (Version: 2.11.3.2) (Waterhouse et al., 2009) using the percentage identity color scheme. Genetic distances were calculated by MEGA (Version: 10.1.8) (Kumar et al., 2018).

## RESULTS

### 3.1 Development of a universal method for dissolving silk from any spider species and its types

To conduct proteome analysis, the method of dissolving spider silk is an inevitable step, which varies depending on the spider species and the types of silk. Previous research explored various techniques, incorporating agents such as LiBr and Gdn-HCl (Geurts et al., 2010; Hu et al., 2005; Kono et al., 2020; Larracas et al., 2016; Pham et al., 2014). Hence, we established a protocol involving 9M LiBr, heat treatment, and sonication, capable of dissolving silk to a certain extent. With this, we successfully extracted proteins from spider silks. The relationship between silk volume and protein content was investigated by dissolving different amounts of silk using a dissolution protocol with LiBr or Gdn-HCl. Subsequently, the protein concentration of the solution was quantified respectively (Fig. 1). Both solvents, LiBr and Gdn-HCl, showed low p-values (1.59e-19 and 6.68e-04), substantiating the correlation between the amount of silk and protein content. Yet, the superior R^2^ values (0.97) associated with LiBr elucidated that this approach extracts protein more effectively from silk compared to Gdn-HCl, which manifested lower R^2^ values (0.24) and a reduced slope (Fig. 1).

**Fig. 1:**
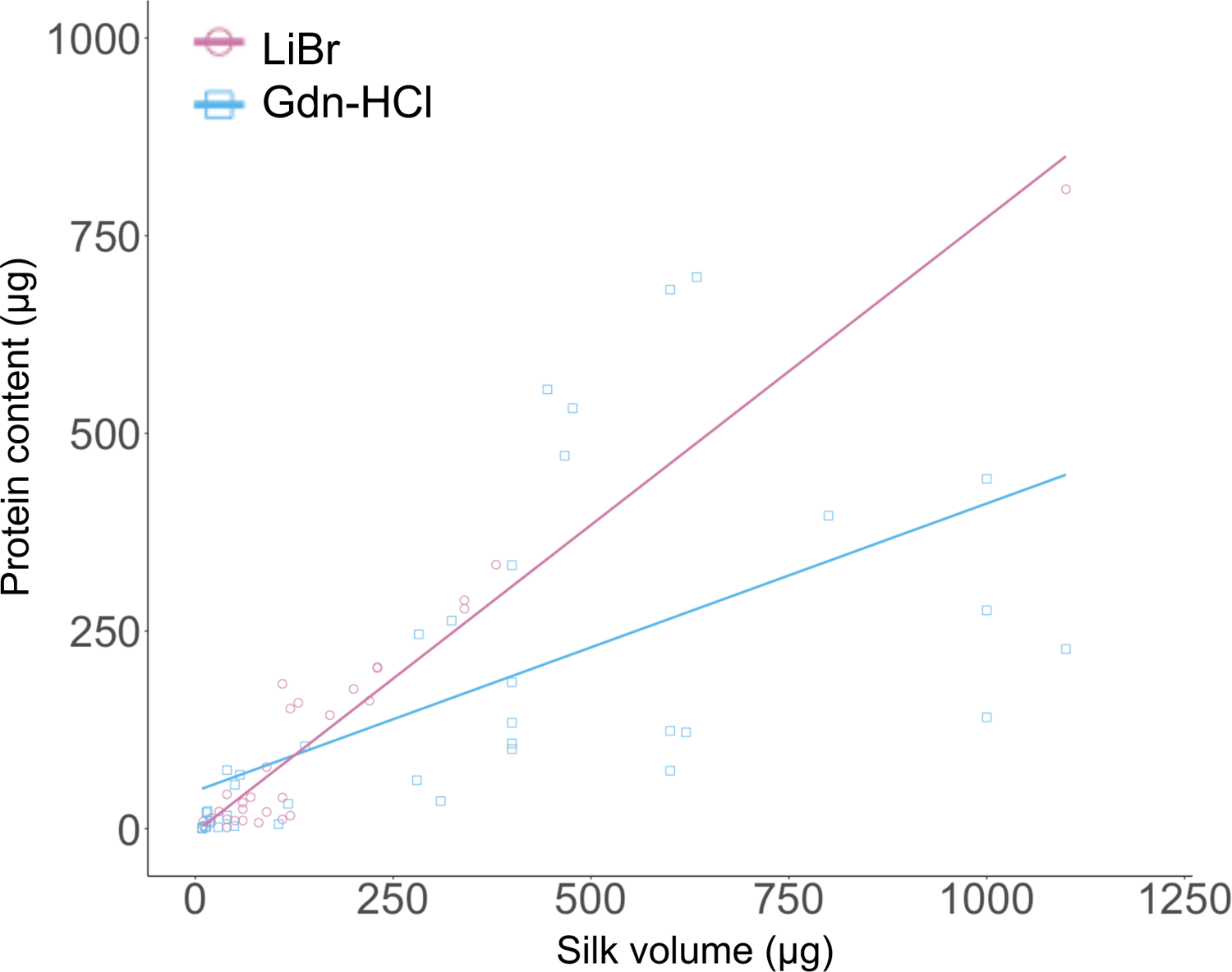
Relationship between silk volume and dissolved protein content. Protein contents were calculated from the protein concentration obtained by BCA quantification. 9M LiBr buffer was used. LiBr: R^2^= 0.97, p = 1.59e-19. Gdn-HCl: R^2^= 0.24, p = 6.68e-04. Silk from the following species was used: *Cyclosa laticauda*, *Cyclosa octotuberculata*, *Dictyna felis*, *Parasteatoda tepidariorum*, *Trichonephila clavata*, *Agelena sylvatica*, *Araneus ventricosus*, *Argiope amoena*, and *Argiope bruennichi*.

### 3.2 Comparison of the accuracy of species estimation

In our proteome analysis, using the list of reference sequences, the spider species was identified through the top-ranking protein in terms of expression and statistical measures, leading us to designate it as the putative species. Subsequently, we investigated whether this putative species matched the silk producer’s species identified through morphology and DNA barcoding. First, we collected webs weaved by captive spiders in a Petri dish, and for those that did not produce webs, we harvested silk using a drill from spiders (Fig. 2ab). Next, we obtained LC-MS/MS data. Given the variety of protein quantification values utilized in proteome analysis, we tested multiple quantification methods. We used two proteome analysis software; PEAKS that performs de novo peptide assembly followed by analysis against the reference gene list (Zhang et al., 2012), and DIA-NN that performs neural network classification from the reference gene list and analyzes MS data (Demichev et al., 2020). These software tools were utilized to calculate iBAQ values (Krey et al., 2014). iBAQ is used to compare and quantify the amount of multiple proteins in one sample, by correcting the MS1 area value by the number of observable peptides which correlates with the length of the protein. As a result, the accuracy of species identification based on iBAQ values was 69.6% with PEAKS, compared to 40% with DIA-NN (Fig. 2c). Next, we calculated pg quantity via DIA-NN, achieving accuracy rate of 42.9% (Fig. 2c). In addition, the accuracy rate was estimated using just MS1 area without any correction for protein length.

**Fig. 2:**
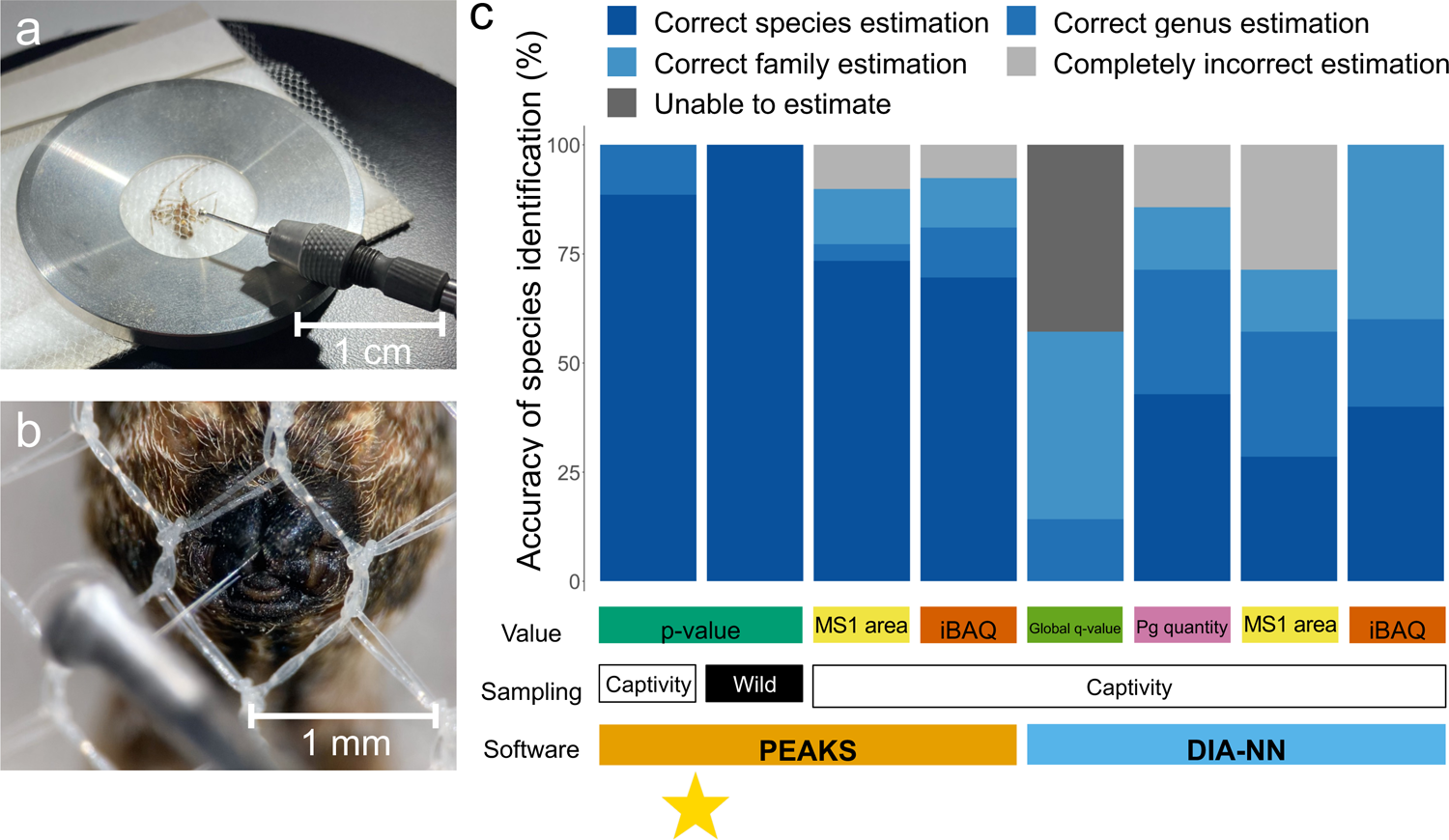
Silk collection and comparison of the accuracy of species estimation in multiple proteome significant difference metrics and quantification values. **a.** Fixation of the spider and positioning of the needle. The spider is fixed with a mesh and the needle attached to the electric drill placed close to the spinnerets. **b.** Enlarged view. The silk was pulled from the spinnerets and wrapped around the needle head using a toothpick. The drill was then switched on and the silk was reeled in. **c.** Comparison of the accuracy of species estimation in multiple proteome significant difference metrics and quantification values. The spider species was estimated based on the top-ranking protein for each value, and it was checked to verify if it matched the true species. The spider species was estimated based on the top-ranking protein for each value, and it was checked to verify if it matched the true species. “Correct genus estimation” if it was correct genus estimation, although incorrect species estimation, “Correct family estimation” if it was correct family estimation, although incorrect genus estimation, “Completely incorrect estimation” if the estimated species belongs to a different family rank. “Unable to estimate” if there are more than two top hits. The indicator with the highest accuracy is marked with a star. For the captive spider webs samples analyzed in PEAKS, n = 79; for the spider webs samples taken from the wild, n = 10; and for the DIA-NN samples, n = 7.

PEAKS achieved a 73.4% accuracy rate, while DIA-NN exhibited an accuracy rate of 28.6% (Fig. 2c). Following this, we extracted the protein that has the highest probability to exist in the sample using statistical significance. PEAKS achieved an 88.6% accuracy rate with p-value. In DIA-NN, the global q-value, which indicates statistical significance, resulted in a 0% accuracy rate. Overall, the comparison of several indicators showed that the strategy of determining the spider species based on the lowest p-value by PEAKS is the most favorable.

### 3.3 Accuracy of species identification in spider webs with contaminants in the field

To evaluate the applicability of this method in field studies, we tested the accuracy of species identification using the same method with spider webs containing contaminants collected from the wild. The presence of contaminants was expected to reduce the overall percentage of spidroin, thereby decreasing the detection of peptides originating from spidroin. Still, we achieved 100% species identification using p-value from PEAKS (n=10) (Fig. 2c).

### 3.4 Limitations of species estimation in silk proteome analysis

We compared the difference of the accuracy rate based on p-value between spider species (Fig. 3a). Interestingly, the accuracy rate of *Araneus ventricosus* (57%) was the lowest compared to other species. Such low accuracy rate occurred when silk derived from *A. ventricosus* were mistakenly identified as MiSp from a congeneric species, *Araneus macacus*. This error was attributed to the high sequence similarity observed in their MiSp N-terminal regions, as illustrated by the alignment result presented in Fig. 3b. Additionally, the genetic distance between those MiSp N-terminus sequences revealed that the genetic distance between *A. macacus* and *A. ventricosus* are closer than those within *A. ventricosus* itself (Fig. 3c). Therefore, it was demonstrated that this method is incapable to distinguish two species with such closely related genetic distances. Despite this limitation, combinations of genetic distances as close as those between *A. ventricosus* and *A. macacus* are relatively infrequent. Out of 4,186 combinations of MiSp N-terminus sequence analyzed from 92 species within the Araneidae family, only 28 combinations exhibited closer genetic distances, as shown in Fig. 3d and Table S1.

**Fig. 3:**
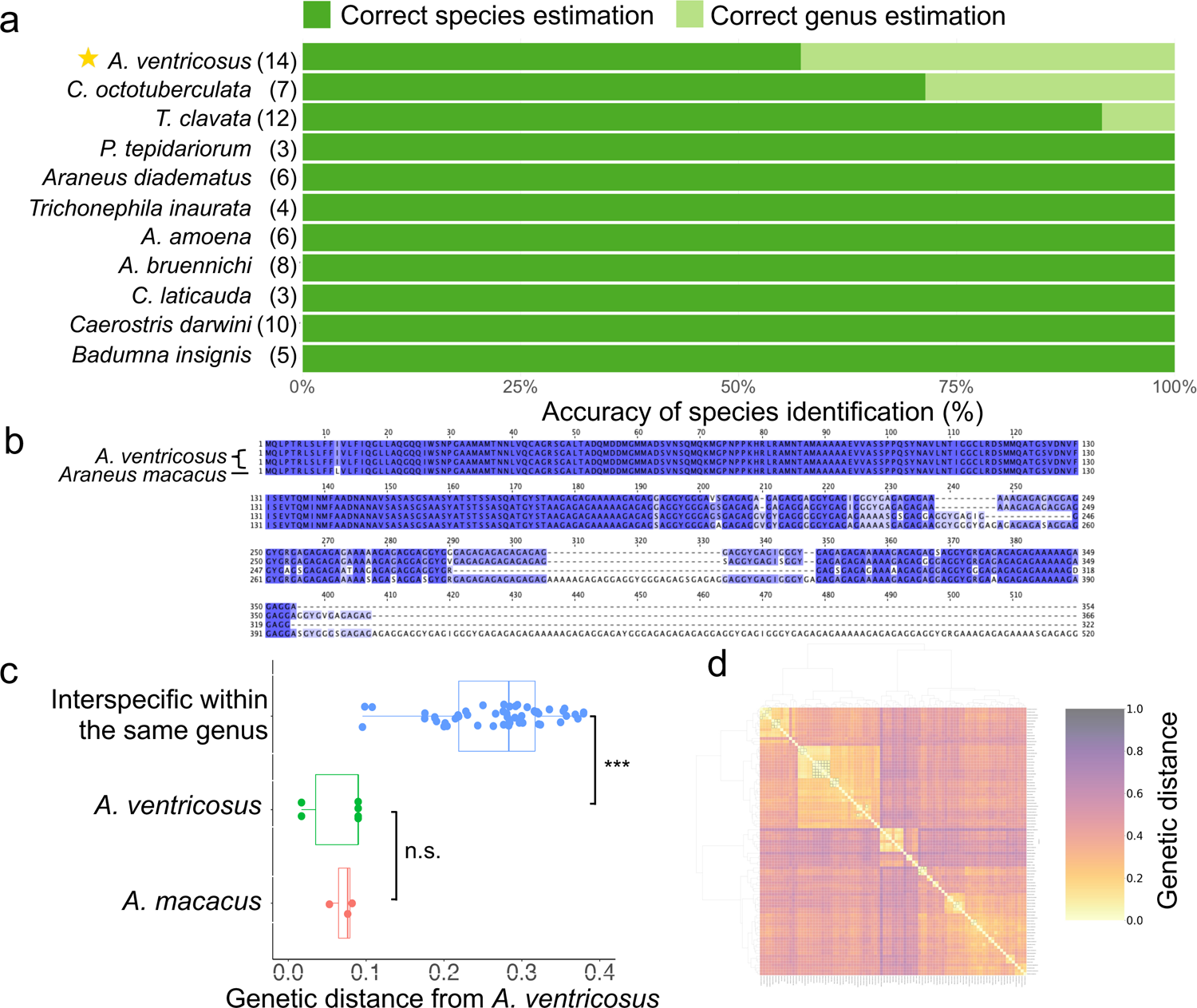
Accuracy and limitations of species estimation in silk proteome analysis. **a.** Accuracy of species estimation by each species using silk proteome analyses based on the smallest p-value obtained by PEAKS. The species with the lowest accuracy (*A. ventricosus*) is indicated with a star. **b.** Alignment of MiSp N-terminus between *A. ventricosus* and *A. macacus*. **c.** Genetic distance between *A. ventricosus* MiSp N-terminus and other MiSp N-terminus, divided into three groups: within *A. ventricosus, A. macacus*, and other species of the genus *Araneus*. Kruskal-Wallis test followed by Dunn’s post-hoc test, adjusted with the Bonferroni method. n.s., non-significant difference (the actual value: 1.000). ***; significant difference: *p*-value < 0.001 (the actual value: 0.000254). **d.** Heat map of the genetic distances among MiSp N-terminus of 92 species within the Araneidae family. Areas with genetic distances smaller than that between *A. ventricosus* and *A. macacus* are outlined in black.

### 3.5 Comparison of the accuracy of species estimation by altering the reference sequence length

Spider silk genes consist of N and C-terminus sequences and repeat sequences in which motifs determine mechanical properties (Ayoub et al., 2007; Gatesy et al., 2001). Due to the presence of repeat sequences, it is difficult to obtain the full-length sequence of spider silk genes, and at present only a few hundred to a few thousand residues from the N and C-terminus have been identified (Fig. S1ab). In addition, the length of spider silk genes currently available varies widely due to differences in the amount of molecular information in the genome and transcriptome of each spider species (Fig. S1ab). Hence, in contrast to the general proteome analyses that use the full length of the gene sequence as a reference, it should be noted that this analysis should consider the fact that the length of the reference gene is partial and varies in length from gene to gene. Therefore, a reference sequence list with only 100 and 200 residues extracted from the N and C-terminus were also prepared and analyzed. Although, the accuracy rate decreased in most proteome indicators, suggesting that the reference gene length should be kept long (Fig. S2).

## DISCUSSION

### 4.1 The possibility of species identification via spider silk proteome analysis

This study displays that it is possible to identify the species by spider silk proteome analysis (Fig. 2c). Here, we demonstrate that species identification is achievable by dissolving spider webs and silks in LiBr, extracting a minimum of 5 μg of protein, and conducting LC-MS/MS mass analysis with DDA using chymotrypsin as the fragmentation enzyme. Identification is feasible through examining the species from which the protein sequences with the smallest p-value originate, as analyzed by PEAKS. Furthermore, if more than 5 μg of protein is extracted, solvents other than LiBr are permissible, and for identification at the family level, employing a different software with DIA poses no issue.

There are some species identification methods that is not dependent on DNA barcoding, and instead protein-based technologies are also available, such as proteome analysis of mammalian bone (Rüther et al., 2022), electrophoresis patterns of animal fur (Sekimoto et al., 2022) and MS patterns of pathogenic bacteria (Holland et al., 1996). Unlike others, our method has a unique feature of being non-invasive and relying on protein sequences. To expand the range of target species, electrophoresis or mass spectrometry must be performed on samples of the target species, as identification methods rely on these experimental data, however, our method requires only peptide sequences as a reference. Therefore, the fact that this experiment demonstrated the ability to identify 15 species, including closely related ones, implies that it is feasible to identify well over 1,000 for which spider silk genes have been collected.

eDNA from spider webs is another non-invasive molecular identification method for spiders (Blake et al., 2016; Xu et al., 2015), which has also been shown to provide biodiversity estimation by identifying organisms attached to the web as an aerial filter (Gregorič et al., 2022). Our method can identify spider species under several conditions: when direct detection of web weavers is necessary, when only a small amount of DNA is available, or when PCR amplification fails. It serves as an alternative to the spider web eDNA method. For example, *Argyrodes* have been known to steal prey by freeloading into the webs of other spider species (Exline & Levi, 1962). Proteome analysis and eDNA of spider webs can reveal both the owner of the web and the spider that freeloaded off it.

In this proteomics analysis study, Data-Dependent Acquisition (DDA) significantly outperformed Data-Independent Acquisition (DIA) results in the scope of species estimation. While DIA, known for its comprehensive analysis capabilities, DDA is considered more suitable for identifying a single most abundant protein with high precision, which was favorable for this method (Bilbao et al., 2015) (Fig. S3).

### 4.2 Future perspective

This method is the first non-invasive molecular biological species identification method that does not harm the organism. In addition, it is widely applicable to the studies of spider biology and ecology and is also useful for the research on wasps parasitizing spiders in the field. For example, by making it possible to identify species from webs without a spider, it is now easier to obtain information about spider habitats from the information provided by their webs than before. Therefore, the World Spider Trait database (Pekár et al., 2021), and the Distribution Data Base of Spiders in Japan (http://www.arachnology.jp/DDBSJ.php) will be greatly expanded and will contribute to the development of spider ecology and the understanding of the population of endangered species. The ability to identify species solely based on spider webs makes research on field spider webs more accessible, it will be a fundamental technology for understanding spider silks, webs, and ecology.

Furthermore, this method is not limited to spider silk, as it can be applied to organisms that release biological materials outside their bodies. For instance, in the case of insects such as the silkworm and the bagworm, it is possible to identify the species by the cocoon and nest structure composed of their silks. In addition, since termites secrete digestive enzymes that break down wood and ticks secrete saliva to their hosts, the proteins contained in their secretions can be used to identify these pests.

### 4.3 Limitation

Although this method is widely applicable, its application is subjected to several limitations. First, this method is not feasible without a protein reference of the target species and are unable to discover new species. Hence, the proteomic analysis results imply the closest species, yet they cannot eliminate the possibility that it is an unlisted spider species in the reference database. This disadvantage, similar to that of DNA barcoding, could see improvements owing to the current exponential expansion of molecular data (Lewin et al., 2018), we can expect it to be gradually augmented and improved.

In addition, it cannot be applied to samples below the limit of measurement. In this study, samples with at least 5 μg of protein were required for preparation, which were enough to detect peptides to estimate species. Also, due to the nature of this method, it cannot be applied to organisms that do not produce biological materials. However, it is expected that future technological innovations in proteome analysis will decrease the minimum limit of measurement and expand the range of applicability of this method, even in organisms that do not produce the purposive protein extended phenotypes, as long as traceable subtle residues are left behind.

## ACKNOWLEDGEMENTS

The authors thank Hiroyuki Nakamura, Maaya Domon, Hironori Iwai, Yu Kurihara, Koh Nakagawa, and Doyoep Kim for their technical support in spider sampling and protein quantification, as well as to Miu Naruki and Kaoru Ogino for their comments on the manuscript. The authors also thank Yuki Baba for enriching discussions. This work was funded by JSPS KAKENHI Grant Number JP21H02210, Osimo Foundation, Taikichiro Mori Memorial Research Fund, and Research funds from the Yamagata Prefectural Government and Tsuruoka City, Japan.

## AUTHOR CONTRIBUTIONS

Conceptualization: N.K., Field Sampling, Silk collection: P.Y. and K.T., Protein extraction, Protein quantification, Peptide purification, Proteome data Analysis, Bioinformatics analysis: P.Y., LC-MS/MS: M.M. and T.M., Writing (original draft): P.Y. and N.K., and Writing (review and editing): All authors.

## DATA AVAILABILITY STATEMENT

The MS raw data and analysis files have been uploaded to the ProteomeXchange Consortium from the jPOST partner repository under accession no. PXD051951.

**Fig. S1: Length distribution of spider silk genes (Spidroin). a.** N-terminus. **b.** C-terminus. 11, 155 spider silk gene sequences of 1, 098 spider species identified in Arakawa *et al*., 2022 [19].

**Fig. S2: Comparison of the accuracy of species estimation in multiple proteome indicators and reference sequence length.** “Correct genus estimation” if it was correct genus estimation, although incorrect species estimation, “Correct family estimation” if it was correct family estimation, although incorrect genus estimation, “Completely incorrect estimation” if the estimated species belongs to a different family rank. “Unable to estimate” if there are more than two top hits. n = 10.

**Fig. S3: Comparison of the accuracy of species estimation in multiple proteome indicators and data acquisition methods.** “Correct genus estimation” if it was correct genus estimation, although incorrect species estimation, “Correct family estimation” if it was correct family estimation, although incorrect genus estimation, “Completely incorrect estimation” if the estimated species belongs to a different family rank. “Unable to estimate” if there are more than two top hits. For the DDA samples, n = 71; for the DIA samples, n = 8.

**Table S1: Combinations of two species within the Araneidae family that are likely indistinguishable with proteome analysis due to the close genetic distance of MiSp N-terminus.** These pairs are emphasized with squares outlined in black in Fig. 3d, exhibiting genetic distances that are closer to or the same as the distance between *A. ventricosus* and *A. macacus*.

