## Supplementary information for "Advancing species identification: A non-invasive molecular approach through spider silk proteome analysis"

Figure S1

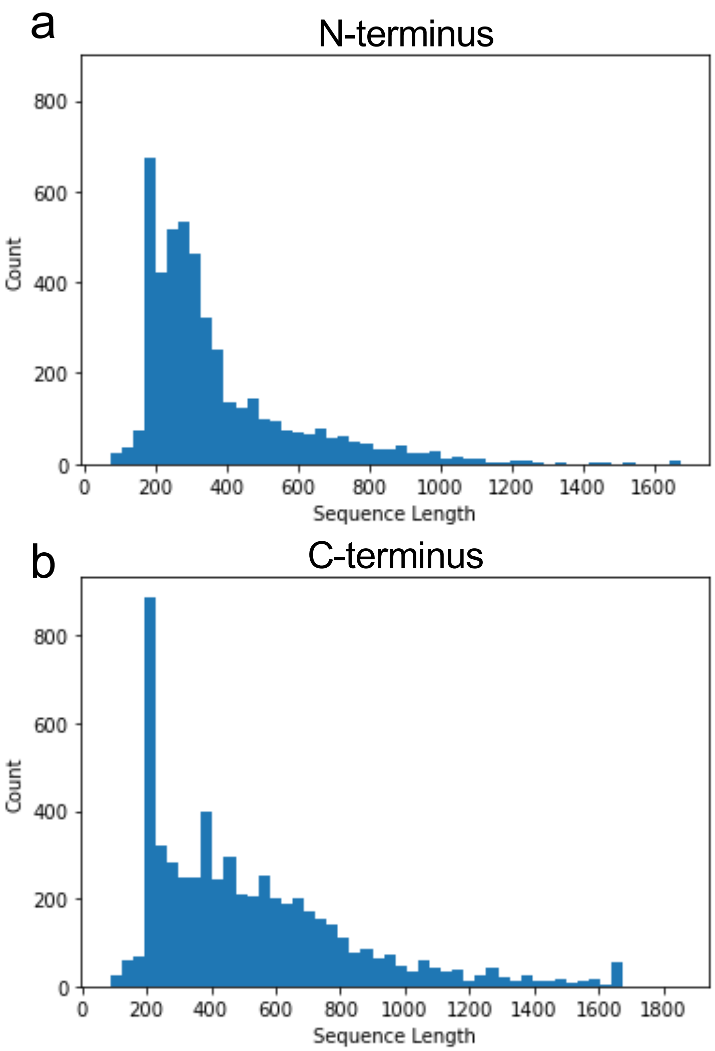
a
N-terminus
Count
Sequence Length
0
200
400
600
800
0
200
400
600
800
1000
1200
1400
1600
b
C-terminus
Count
Sequence Length
0
200
400
600
800
0
200
400
600
800
1000
1200
1400
1600
1800

Figure S2

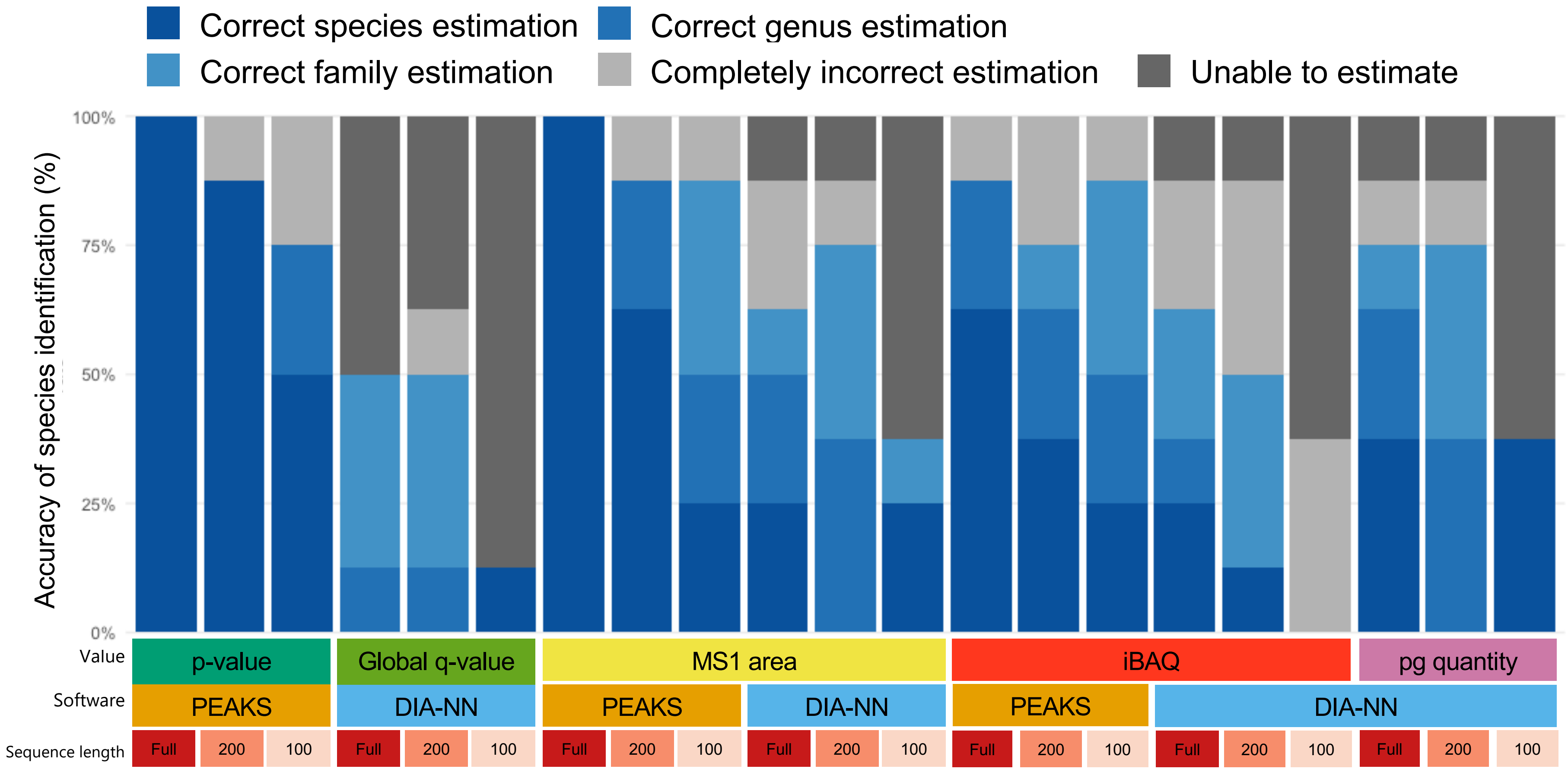

Correct species estimation
Correct genus estimation
Correct family estimation
Completely incorrect estimation
Unable to estimate
Accuracy of species identification (%)
100%
75%
50%
25%
0%
Value
Software
Sequence length
p-value
Global q-value
MS1 area
iBAQ
pg quantity
PEAKS
DIA-NN
PEAKS
DIA-NN
PEAKS
DIA-NN
Full
200
100

Figure S3

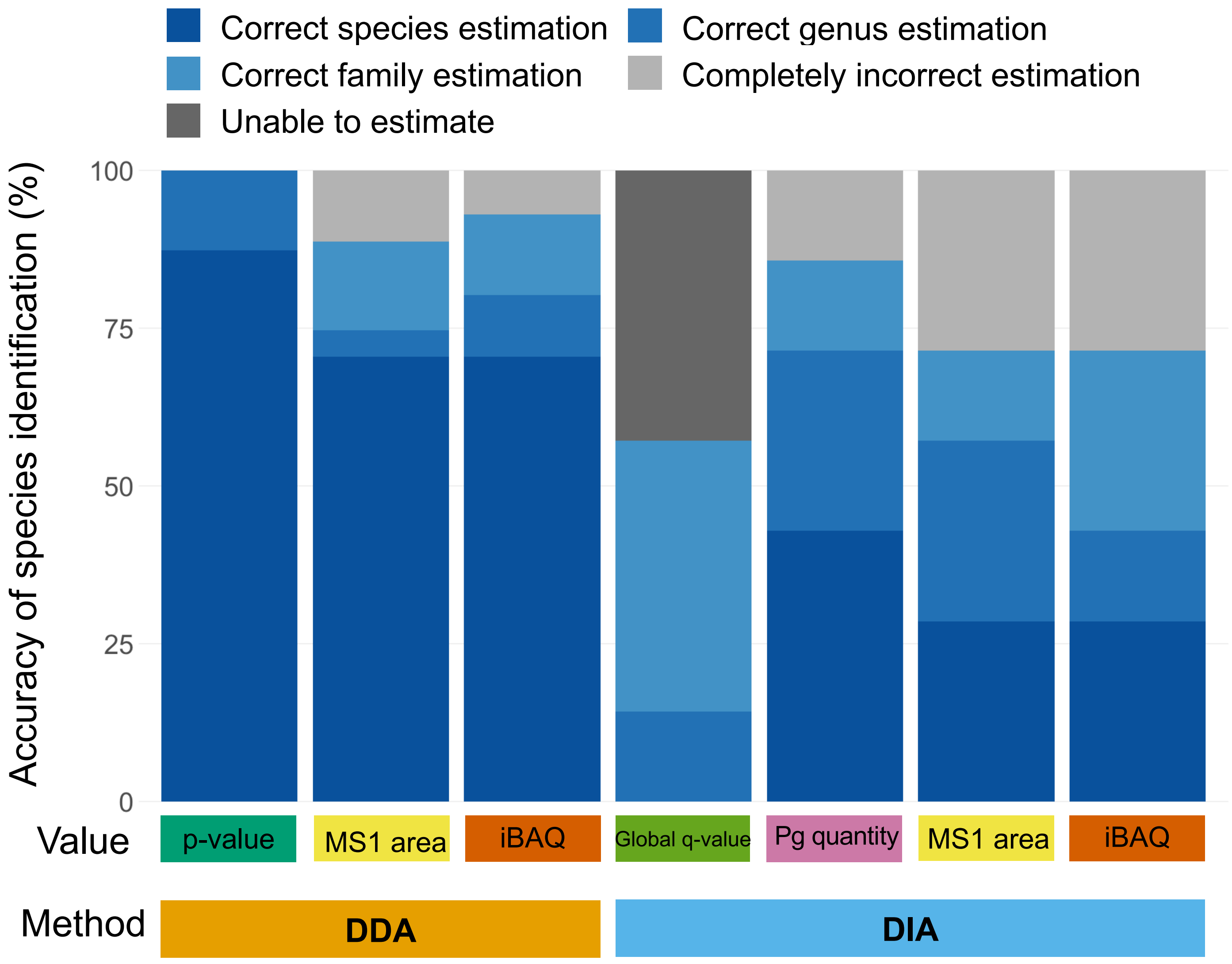

Correct species estimation
Correct genus estimation
Correct family estimation
Completely incorrect estimation
Unable to estimate
Accuracy of species identification (%)
100
75
50
25
0
Value
p-value
MS1 area
iBAQ
Global q-value
Pg quantity
MS1 area
iBAQ
Method
DDA
DIA

Table S1

| First | Second | Genetic distance |
|---|---|---|
| *Neoscona mellotteei* | *Neoscona scylloides* | 0.050 |
| *Araneus bicentenarius* | *Araneus seminiger* | 0.056 |
| *Araneus seminiger* | *Araneus uyemurai* | 0.057 |
| *Araneus macacus* | *Araneus ventricosus* | 0.062 |
| *Argiope aemula* | *Argiope boesenbergi* | 0.056 |
| *Argiope aemula* | *Argiope keyserlingi* | 0.052 |
| *Argiope boesenbergi* | *Argiope keyserlingi* | 0.062 |
| *Cyrtophora exanthematica* | *Cyrtophora unicolor* | 0.055 |
| *Cyrtophora ikomosanensis* | *Cyrtophora moluccensis* | 0.044 |
| *Plebs eburnus* | *Plebs sachalinensis* | 0.056 |
| *Plebs eburnus* | *Plebs yanbaruensis* | 0.052 |
| *Plebs sachalinensis* | *Plebs yanbaruensis* | 0.026 |
| *Cyclosa argenteoalba* | *Cyclosa kumadai* | 0.040 |
| *Cyclosa argenteoalba* | *Cyclosa confusa* | 0.036 |
| *Cyclosa argenteoalba* | *Cyclosa japonica* | 0.029 |
| *Cyclosa argenteoalba* | *Cyclosa monticola* | 0.046 |
| *Cyclosa kumadai* | *Cyclosa confusa* | 0.048 |
| *Cyclosa kumadai* | *Cyclosa japonica* | 0.044 |
| *Cyclosa kumadai* | *Cyclosa monticola* | 0.053 |
| *Cyclosa kumadai* | *Cyclosa omonaga* | 0.056 |
| *Cyclosa confusa* | *Cyclosa japonica* | 0.006 |
| *Cyclosa confusa* | *Cyclosa monticola* | 0.011 |
| *Cyclosa confusa* | *Cyclosa omonaga* | 0.032 |
| *Cyclosa japonica* | *Cyclosa monticola* | 0.011 |
| *Cyclosa japonica* | *Cyclosa omonaga* | 0.024 |
| *Cyclosa monticola* | *Cyclosa omonaga* | 0.040 |
| *Cyclosa laticauda* | *Cyclosa octotuberculata* | 0.028 |
| *Cyclosa laticauda* | *Cyclosa onoi* | 0.045 |
| *Gasteracantha cancriformis* | *Gasteracantha diadesmia* | 0.062 |
